# Efficient and flexible strategies for gene cloning and vector construction using overlap PCR

**DOI:** 10.1101/101147

**Authors:** Zhongtian Liu, Tingting Zhang, Kun Xu

## Abstract

Gene cloning and vector construction are basic technologies in modern molecular biology for gene functional study. Here, we present flexible and efficient strategies for gene cloning and vector construction using overlap PCR. We firstly cloned the open reading frames (ORFs) of the porcine *MSTN*, chicken *OVA* and human *α-glucosidase* genes by overlap PCR-based assembling of their exons, which could be amplified with genomic DNAs as the templates without RNA extraction and RT-PCR reaction. Secondly, we generated additionally three designed functional cassettes by overlap PCR-based assembling of different DNA elements, which facilitated the construction their expression vectors greatly. Moreover, we further developed an interesting overlap-circled PCR method for fast plasmid vector construction without any cutting and ligating procedure. These advanced applications of overlap PCR provide useful alternative tools for gene cloning and vector construction.

## Introduction

Gene cloning and vector construction are the foundational and routine technologies for molecular biological research. The first step for studying a particular gene is usually to clone it from the cDNA using RT-PCR, and routinely the next step is to construct corresponding expression vector by restrictive endonuclease-based digestion and ligase-based ligation. The restrictive endonuclease-ligase dependent strategy for gene cloning has been deeply developed and widely used. However, there are inherently several limitations, such as the requirement of high-qualified cDNA template for desired gene cloning, the inconvenience for simultaneous multiple gene cloning and the limited restriction enzyme sites available on the backbone plasmid for vector construction. Multiple fragment cloning can be achieved by enzymatic assembly of overlapping DNA fragments with the alternative application of 5ʹ-exonucleases [1, 2] and by Golden Gate cloning using type IIs restrictive enzymes [3-5].

Besides, a series of nuclease or/and ligase-independent strategies for constructing vectors based on different characteristics of intent terminal sequences has been developed, such as the enzyme-free cloning using annealing tails [6], the T-A cloning applying special ends of PCR products[7], the UDG or USER cloning with Uracil DNA glycosylase [8, 9], the Gateway cloning dependent on bacterial homologous recombination (HR) [10], the mating-assisted genetically integrated cloning (MAGIC) using bacterial *in vivo* site-specific endonuclease cleavage and HR [11], and the sequence and ligation-independent cloning (SLIC) using *in vitro* HR and single-strand annealing (SSA) [12]. Although these methods mentioned above all have their own applications and advantages, new improvements leading to more flexible, feasible, convenient and economical approaches for gene cloning and vector construction are still needed.

Here, we demonstrated that the open reading frame (ORFs) of functional genes could be assembled with exon amplicons from genomic DNA using overlap PCR strategy, which is useful and practical for amplifying long eukaryotic genes interrupted by introns [13-17]. Besides, multiple genes or DNA elements are sometimes required to be cloned into a single plasmid vector, which is always inconvenient with multiple cloning steps and confined by limited cloning sites available. The overlap PCR strategy could be effective for assembling these genes or elements. We further demonstrated that the overlap-circled PCR could be used to assemble the intent DNA fragment and the plasmid backbone for vector construction.

## Results

### Cloning eukaryotic genes by overlap PCR-based assembling of their exons

Firstly, we tried to clone the ORF of porcine *MSTN* gene (NC_010457) by the overlap PCR strategy. Myostatin is a myokine protein released by myocytes to inhibit myogenesis and the *MSTN* gene contains 3 exons and 3 introns (Fig.1A). Generally, to obtain the 1128 bp CDS, the exons were amplified respectively via PCR with porcine genomic DNA as template and were assembled one by one via overlap PCR. The principle for designing overlap PCR primers is illustrated in Fig.1B. The 3 exons were amplified with primer pairs of exon 1-F/exon 1-R, exon 2-F/exon 2-R, and exon 3-F/exon 3-R respectively (Fig.1B and 1D). The PCR products of these 3 exons were purified by gel extraction, and further served as the templates for generating the whole ORF fragment of *MSTN* gene (Fig.1C) by the second overlap PCR step (Supplementary Table S2 and Fig.S2) with primers exon 1-F and exon 3-R (Fig.1B and 1E). This technique makes gene cloning simply by exon-ligation through overlap PCR without RNA extraction and RT-PCR, and can be applied widely for amplifying various eukaryotic gene interrupted by introns for molecular biology study.

**Figure1.**
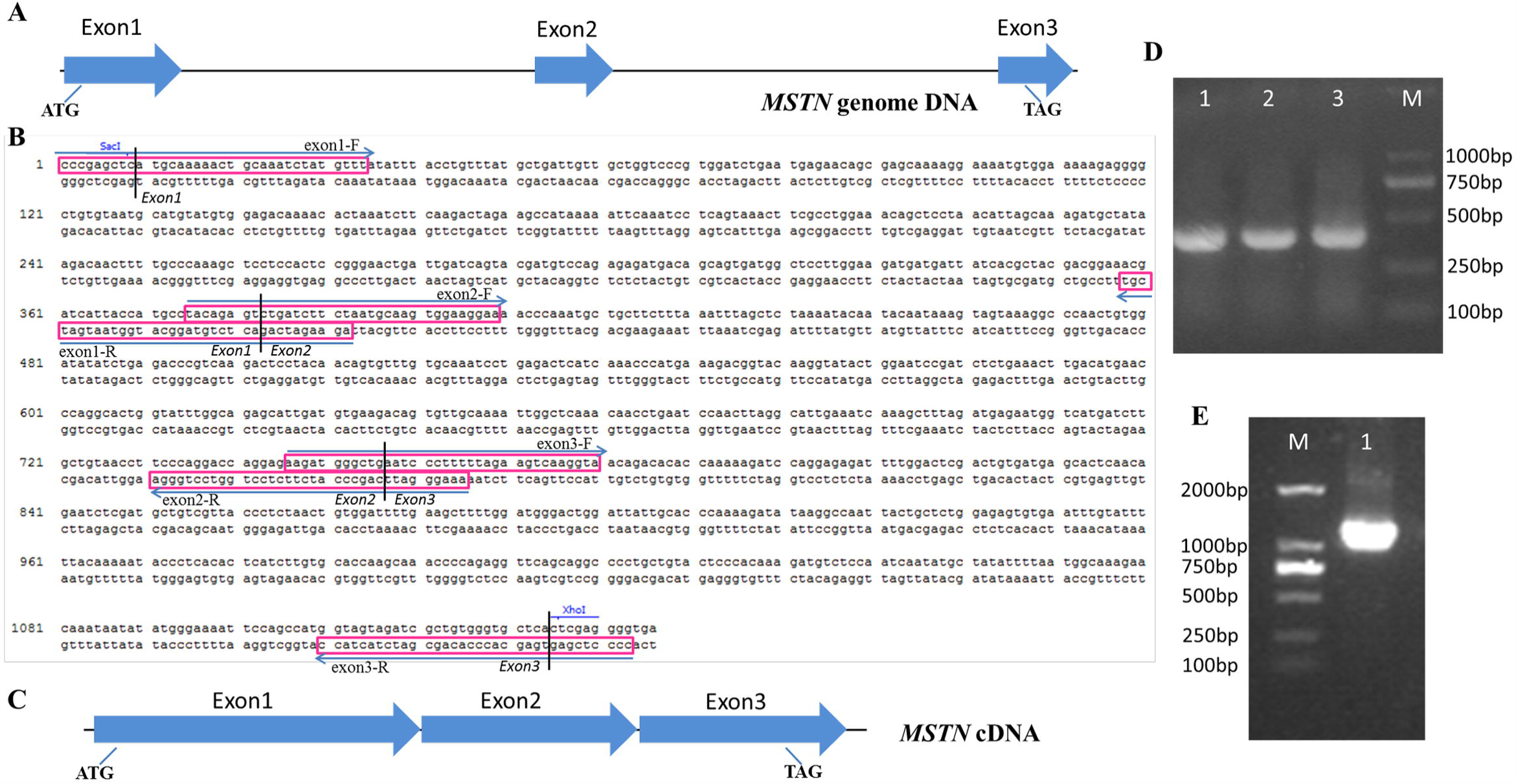
Cloning the porcine *MSTN* gene by assembling of exons. **A.** The schematic diagram of porcine *MSTN* gene in genome. The *MSTN* gene contains three exons, which are illustrated by solid arrows in blue. The start codon in exon 1 and stop codon in exon 3 labeled. **B.** The design of the overlapping primers. Primers are boxed and their orientations are indicated by parallel arrows. Exon-exon junctions are marked by vertical lines. The genome DNA sequence for the *MSTN* gene is partially displayed. **C.** Schematic illustration of the *MSTN* cDNA generated by assembling of exons. **D.** Agarose gel electrophoresis detection for three PCR amplified exon fragments of *MSTN*. Lane M: DL200 marker, lane 1, 2, 3: exon 1, exon 2 and exon 3. **E.** Agarose gel electrophoresis detection for the overlapped *MSTN* cDNA fragment. Lane M: DL200 marker, lane 1: overlapped *MSTN*.

To demonstrate the feasibility, the chicken ovalbumin gene (NM205152, *OVA*) containing 8 interval exons (with the codon ATG in exon 2) inlayed in the 7573 bp genomic DNA was assembled similarly. Seven pairs of overlap PCR primers were designed as described above. The 1161 bp ORF of *OVA* gene was generated by assembling fragments from exon 2 to exon 8 (Supplementary Fig.S1A and S1B). We next successfully assembled the 2132 bp ORF of human *α-glucosidase* gene with 11 exons (Supplementary Fig.S1C and S1D). Actually, in addition to the two-step overlap PCR procedure described above, the intent ORF fragments of the two genes could also obtained by one-step overlap PCR simply pooling all the overlap PCR primers, the genomic DNA template and the PCR regents together in a single tube. These studies suggested that the overlap PCR strategy is effective and practical for amplifying long eukaryotic genes interrupted by introns, which may be more convenient and economical when applied in the further studies.

### Cloning functional cassettes by overlap PCR-based assembling of different DNA elements

In order to further verify the advantage of the overlap PCR strategy, we tried to assemble different DNA elements rapidly to generate designed functional cassettes. Firstly, the SV40P.neo-IRES-tk-polyA cassette was designed and constructed. Three pairs of overlapping primer were designed as shown in Fig.2A. The SV40P.neo, IRES and tk-polyA fragments were amplified from plasmid DNA through touch-down PCR [18] and purified by gel extraction routinely. The whole 3.3 kb fragment was generated by overlap PCR with primers at the ends using the three fragments as the templates (Fig.2B).

**Figure2.**
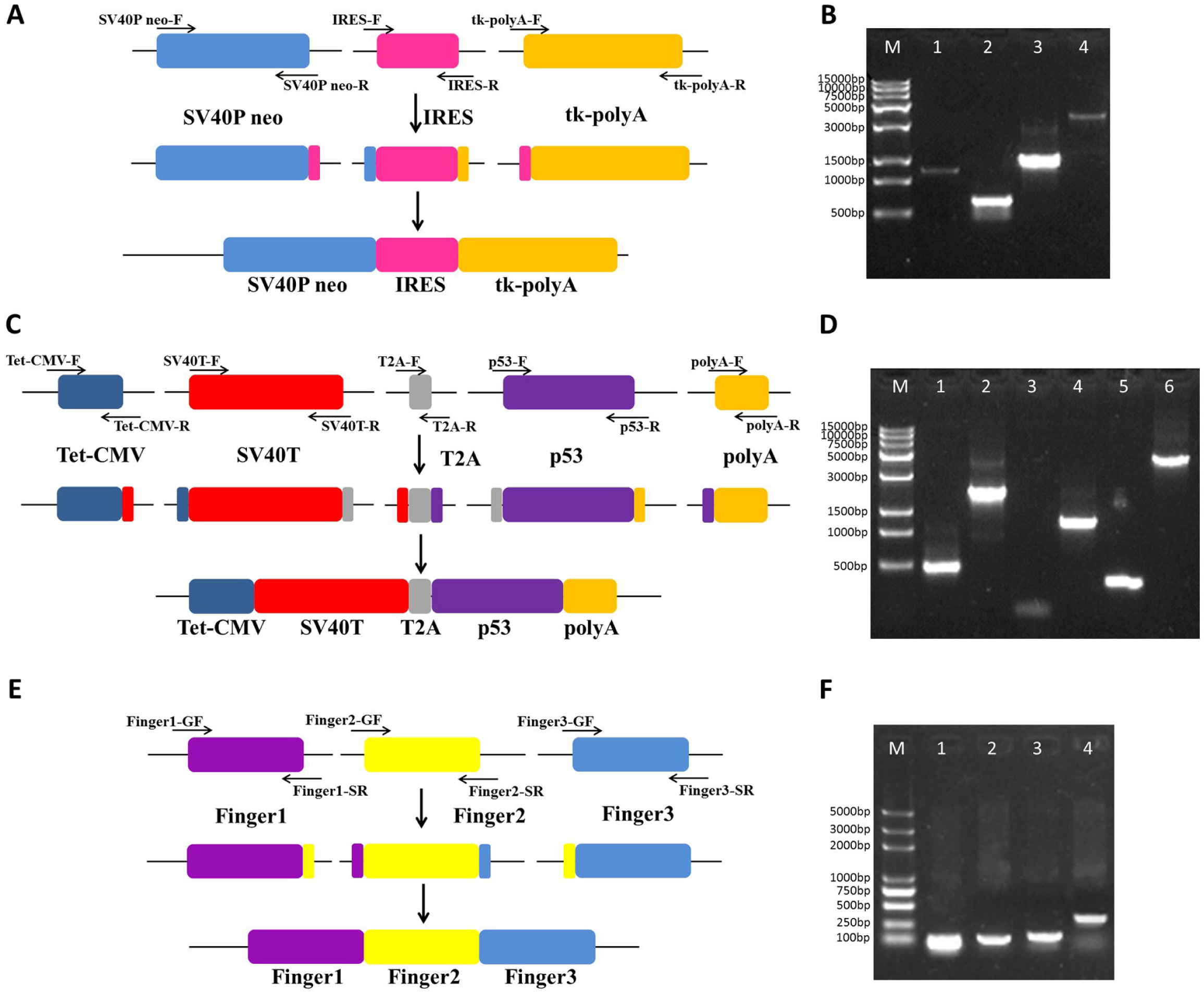
Cloning functional cassettes by assembling of different DNA elements. **A.** Schematic diagram for assembling the SV40P.neo-IRES-tk-polyA cassette. **B.** Agarose gel electrophoresis detection of the PCR products. Lane M: Trans 15K marker, lane 1, 2, 3, 4: represent respectively the PCR products of SV40P. neo, IRES, tk-polyA and overlappedSV40P.neo-IRES-tk-polyA. **C.** Schematic diagram for assembling the Tet.SV40T-T2A-p53-polyA cassette. **D.** Agarose gel electrophoresis detection of the PCR products. Lane 1, 2, 3, 4, 5, 6: Tet.CMV, SV40T, T2A, p53, polyA and overlapped Tet.SV40T-T2A-p53-polyA. **E.** Schematic diagram for assembling the zinc finger cassette. F. Agarose gel electrophoresis detection of the PCR products. Lane M: Trans 2K marker, lane 1, 2, 3, 4: Finger 1, Finger 2, Finger and fused zinc fingers. All overlapping primers designed were labeled and indicated by arrows.

Secondly, we tried to assemble a larger Tet.SV40T-T2A-p53-polyA cassette, which was intended to be 5.7 kb in length. Five pairs of overlapping primers were designed as shown in Fig.2C. Corresponding DNA elements were PCR amplified from either plasmid DNA or genomic DNA, and were assembled together successfully by overlap PCR (Fig.2D). Furthermore, by using the overlap PCR strategy, we also obtained the zinc finger cassette with three fingers for targeting the *CCR5* gene (Fig.2E and 2F). The successful assembly of different DNA elements through overlap PCR will facilitate the fast cloning of functional cassette by avoiding multiple cloning steps.

### Fast plasmid vector construction by overlap-circled PCR

In another attempt, we obtained a circle plasmid containing the selective marker gene and the *MSTN* gene expression cassette through overlap-circled PCR (Fig.3). To begin with, the backbone plasmid pBlueScript (Addgene) containing the *Amp^r^* gene and pUC replication initiation site was amplified with primers pBlue-F/pBlue-R. Then the ORF of eukaryotic *MSTN* gene was cloned by the two-step overlap PCR strategy as described above. Finally, the pBlueScript backbone fragment and the *MSTN* ORF fragment were overlapped by circled PCR [14, 15] using the *pfu* DNA polymerase without primers for generating the pBlueScript-MSTN construct, which was used to transform the *E.coli* JM109 competent cells for the amplification. Then plasmid DNA was extracted from culture of positive colonies, the digesting assay with *Eco*R I/*Xho* I restrictive enzymes was conducted and agarose gel electrophoresis was performed to confirm the positive plasmid clones. The results revealed that 4 out of the 12 clones (33.3%) were positive. Here, we demonstrated a useful overlap-circled PCR method for fast vector construction, through only two or three PCR procedures without using any ligase and endonuclease.

**Figure3.**
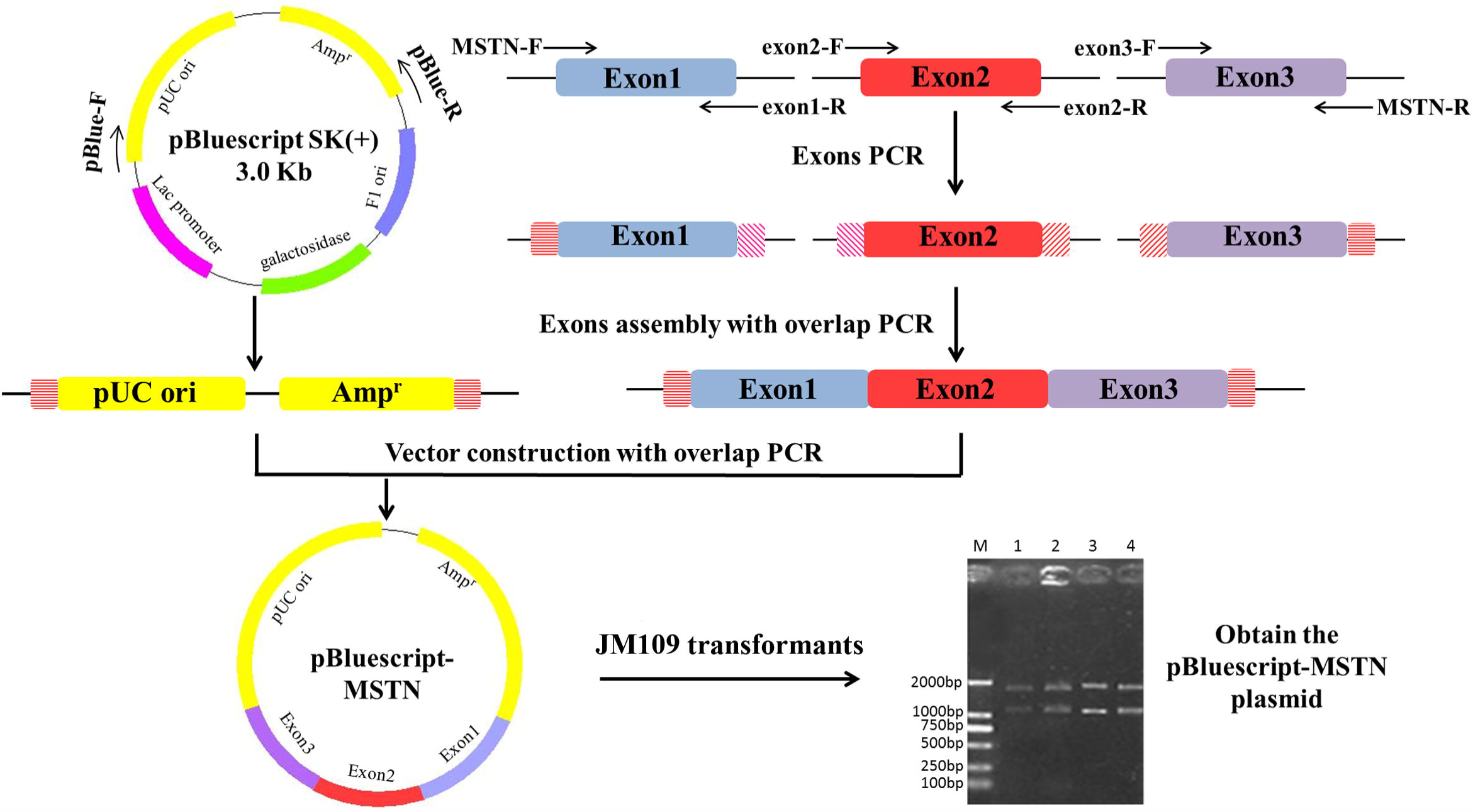
Schematic illustration for the overlap-circled PCR-based plasmid vector construction. The *Amp^r^* marker gene and the pUC replication initiation site of pBlueScript backbone were amplified by normal PCR and the MSTN fragment was generated by overlap PCR as described above. The pBlueScript backbone fragment and the MSTN fragment were overlapped by circled PCR using the pfu DNA polymerase without primers for generating the pBlueScript-MSTN construct. *E.coli* JM109 were transformed with the product for cloning and the gel represents the result of digesting assay with *Eco*R I/*Xho* I. Lane M: DL200 marker; lane 1-4: plasmids extracted from positive JM109 transformants; the two intent bands should be 1168 bp and 1776 bp respectively in length.

## Discussion

In summary, we firstly cloned the ORFs of genes porcine *MSTN*, chicken *OVA* and human *α-glucosidase* by overlap PCR-based assembling their exons. Secondly, we successfully obtained several functional cassettes by assembling different DNA elements from plasmid or genomic DNA. Thus, the first advantage of the overlap PCR strategy is that it is much convenient and cost-effective to conduct gene cloning without RNA extraction and RT-PCR, and the cloning can be even achieved by one step overlap PCR procedure. The second advantage is that functional cassettes can be generated by quickly assembling corresponding DNA elements, facilitating the fast cloning of complicated cassettes by avoiding multiple cloning steps. Finally, we further demonstrated an interesting overlap-circled PCR method for fast vector construction with only PCR procedures, which makes the vector construction much easier and more economic by simply designing several pairs of primers without using any ligase and endonuclease. To sum up, our work demonstrated that the overlap PCR strategy can be flexible, effective, convenient and economical for gene cloning and vector construction.

## Materials and Methods

### Chemicals and stains

*Taq* DNA polymerase, *Pfu* DNA polymerase, DNA marker, *E.coli* JM109 competent cells were purchased from TransGen Biotech (Beijing, China). T4 DNA ligase and restriction endonucleases were from NEB (Beijing, China).

### Preparation of genomic DNA

The genome DNAs prepared from tissues (pig and chicken) was shown as follows and that from cultured cells (human 293T) was manipulated similarly without the digestion. Digest approximately 2 mm^3^ tissues with 300 μL digestion buffer [5 mM EDTA, pH 8.0, 200 mM NaCl, 100 mM Tris, pH 8.0, 0.2% sodium dodecyl sulfate (SDS)] with 0.4 mg proteinase K/1 mL digestion buffer, in a 1.5 mL tube at 55 °C overnight. Add an equal volume (about 300 μL) of phenol/chloroform to each sample, and vortex for 30 seconds. Centrifuge at room temperature at max speed for 5 minutes to separate phases. Transfer the upper phase of each sample to a fresh tube. Add 1 mL 100% ethanol into each tube, mix completely. Centrifuge at max speed for 10 minutes, and then pour out the ethanol. Wash the DNA pellets by adding 1 mL 70% ethanol into the tubes, then centrifuge at max speed for 10 minutes. Pour out the ethanol from the tubes, immediately dissolve the DNA pellets with 50 μL TE buffer (10 mM Tris-HCl, 0.2 mM Na_2_EDTA, pH 7.5) after air-drying.

### Overlap PCR for assembling exons and DNA fragments

The individual exons and DNA element fragments were amplified through standard PCR procedure and reaction system (15μL) as shown in Supplementary Table2. The overlap PCR for assembling exons and different DNA fragments were performed in 50μL reaction system (Supplementary Table2) using the touch-down PCR procedure (Supplementary Fig.S2). Briefly, the touch-down PCR parameters used were as follows: pre-denaturation at 95°C for 5 minutes; followed by 18 touch-down cycles of denaturation at 95°C for 30 seconds, annealing at 68°C for 30 seconds with one degree reduction in each cycle, and polymerization at 72°C for 30 seconds; then, 25 cycles of denaturation at 95°C for 30 seconds, annealing at 50°C for 30 seconds, and polymerization at 72°C for 30 seconds; and a final incubation for 10 minutes at 72°C; end with 10°C incubation. PCR products were purified by gel extraction using the kit according to the manufacturer’s instructions (OMEGA; Guangzhou, China). All primers used in this work were ordered from Genscript (Nanjing, China) and the sequence information was shown in Supplementary Table1.

### Vector construction by overlap-circled PCR

For the plasmid vector construction by overlap-circled PCR, the plasmid backbone and the insertion fragments with overlapping ends were firstly obtained by respectively normal and overlap PCR amplification. Then, the two fragments were overlapped by circled PCR [14, 15] using the *pfu* DNA polymerase without primers for generating the intent construct. The product was directly used to transform *E.coli* JM109 competent cells. Positive colonies were picked and cultured for plasmid DNA extraction using the kit (OMEGA; Guangzhou, China). The digesting assay with *Eco*R I/*Xho* I restrictive enzymes and agarose gel electrophoresis was conducted to confirm the positive plasmid clones.

## Acknowledgements

The authors would like to thank all the colleagues in Professor Zhang’s lab for their excellent technical assistance and helpful discussions.

## Author contributions

Zhongtian Liu and Tingting Zhang performed the experiments; Kun Xu designed the experiments and wrote the manuscript; Tingting Zhang and Kun Xu funded this work.

